# B-Cell Involvement in Immune Checkpoint Inhibitor-Induced Lichen Planus: A Comparative Analysis with Non-Drug-Related Lichen Planus

**DOI:** 10.1101/2024.01.04.574049

**Authors:** Alice Tison, Delphine Legoupil, Marion Le Rochais, Patrice Hémon, Nathan Foulquier, Quentin Hardy, Sophie Hillion, Arnaud Uguen, Jacques-Olivier Pers, Laurent Misery, Divi Cornec, Soizic Garaud

**Author notes:** **Corresponding author:** Soizic Garaud, PhD 9 Rue Félix le Dantec 29200 Brest, France. These authors contributed equally to the present work (Alice Tison, Delphine Legoupil).

## Abstract

**Background:** Immune checkpoint inhibitors (ICI) have significantly improved cancer prognosis but can lead to immune-related adverse events (irAE), including cutaneous manifestations affecting 30% to 60% of ICI-treated patients. However, the physiopathology of cutaneaous irAE remains unclear.

**Objective:** This study investigated the immune infiltration in tissues affected by cutaneous irAE to elucidate their contribution to the pathogenesis of these toxicities.

**Methods:** Skin biopsies from 6 patients with ICI-induced lichenoid eruptions were compared using imaging mass cytometry to samples from 7 controls with non-drug-related lichen planus.

**Results:** T cells were the predominant cell type within the inflammatory infiltrate in all samples, but we observed a reduced T-cell infiltration and an increased B-cell frequency in ICI-induced lichen planus compared to non-drug related lichen planus. Among B cells, we observed a significant decrease in IgD-CD27-double-negative B cells and an increase in IgD+CD27-naïve B cells. Spatial analysis demonstrated that infiltrating B cells were organized in aggregates close to T cells in ICI-induced lichen planus.

**Limitations:** This is a retrospective single-center study with a relatively small sample size.

**Conclusion:** This study sheds light on the involvement of B cells in the pathogenesis of ICI-induced lichen planus, suggesting distinct immunological mechanisms from non-drug-related lichen planus.

**CAPSULE SUMMARY:**

- Lichenoid manifestations are a common but understudied side effect occurring in patients receiving anti-PD-1 antibodies.
- ICI-induced lichen planus displays distinct physiopathology from non-drug-related lichen planus, with a decrease of T-cell infiltration concomitantly to the increase of B cells organized in aggregates.

## BODY OF MANUSCRIPT

Immune checkpoint inhibitors (ICI) have significantly improved cancer outcomes; however, the release of immune regulatory controls can induce inflammatory side effects known as immune-related adverse events (irAE) affecting multiple organs. One common type of IrAE is cutaneous irAE (c-irAE) which occurs in approximately 30% to 60% of patients receiving ICI.^1-3^ They have a wide phenotypic range, including eczematous maculopapular rashes, pruritus, lichenoid reactions, psoriasis, acneiform rashes, vitiligo-like lesions, autoimmune skin diseases (e.g., bullous pemphigoid, dermatomyositis, alopecia areata), sarcoidosis or nail and oral mucosal changes, and Stevens-Johnson syndrome/toxic epidermal necrolysis.^4,5^ Moreover, the use of anti-CTLA-4 and anti-PD-1 therapies in combination is associated with the development of more frequent, more severe and earlier c-irAE compared to single agents.^6^ While a majority of dermatologic toxicities are self-limiting and readily manageable, it is not uncommon that they result in severe or irreversible skin involvement, impairment of patients’ quality of life and potentially life-threatening consequences.

The exact immunological mechanisms responsible for c-irAE remains unclear. However, several case reports of lichenoid reactions induced by ICI (ICI-LP) have demonstrated T-cell infiltrates with a predominance of helper T-cells over cytotoxic T-cells, with approximately 10% of the T-cells staining positive for PD-1 and a greater histiocytic infiltrates compared to treatment-naive lichen planus.^7-9^ Moreover, a retrospective analysis of melanoma patients with secondary cutaneous maculopapular eruptions revealed a predominance of T cells with isolated B cells.^10^ A small subpopulation of regulatory T-cells was also identified in the dermis mainly with a perivascular distribution.

Our study aimed to provide a comprehensive analysis of immune infiltration in ICI-LP, in comparison with non-drug related lichen planus (LP), using single-cell high dimensional imaging mass cytometry (IMC). We highlighted the reduction of T-cell infiltration in ICI-LP, concomitantly to an increased frequency of B cells. Phenotypic characterization of B-cell infiltrates revealed a significant decrease in IgD-CD27-double negative B cells in association with an increase in IgD+CD27-naïve B cells in ICI-LP compared to treatment-naïve patients. Finally, spatial analysis demonstrated that infiltrating B cells form aggregates close to T cells in ICI-induced lichen planus. Overall, this study provides insights into the role of B cells in the pathogenesis of ICI-LP relevant for the development of diagnostic biomarkers and the development of novel therapeutic strategies.

## METHODS

### Patient population and clinical samples

This retrospective single-center study included patients who developed lichenoid reactions after receiving pembrolizumab and referred to the Department of Dermatology at Brest Hospital between January 2020 and December 2022. Punch biopsies specimen from these patients were formalin-fixed and paraffin-embedded (FFPE) and were compared with biopsies of LP used as a control group.

All the studies involving human tissues were approved by the local institutional ethics committee.

### Hematoxylin and eosin staining

Skin biopsy specimens were fixed in 10% buffered formalin, processed, and embedded in paraffin using standard histopathologic methods in the department of pathology. Four µm sections were stained with hematoxylin and eosin (H&E). Stained sections were scanned and visualized using the NanoZoomer-SQ digital slide scanner (Hamamatsu) and its viewing platform (NDP.Viewer).

### Imaging Mass Cytometry staining and acquisition

We designed an IMC panel consisting of twelve metal-tagged antibodies, covering both non-immune and immune cells to evaluate the tissue microenvironment of skin biopsies (Table 1). A nuclear intercalator dye (Iridium) was included to allow the identification and segmentation of individual cells. All antibodies used for IMC were validated by immunohistochemistry for specific staining patterns.

**Table 1.**
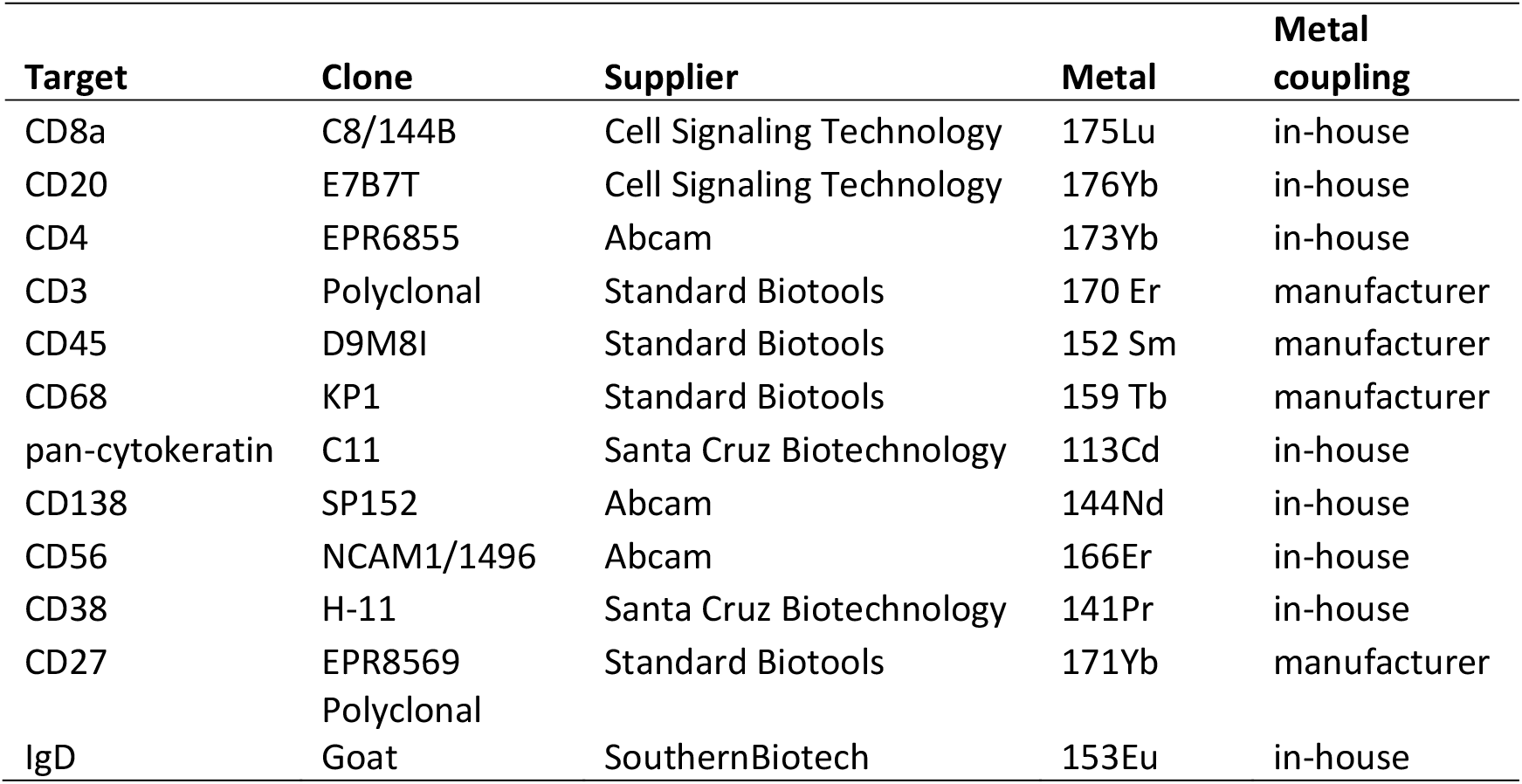
Antibodies used for IMC.

Some antibodies were conjugated to metal tags using the Maxpar® antibody labeling kit (Standard BioTools) following the manufacturer’s instructions. The yield of each conjugation was determined by using the NanoDrop spectrophotometer, and antibodies were stored at a final concentration of 0.5 mg/mL in stabilizing PBS (Candor Biosciences) at 4°C. All antibodies were titrated on tonsil section by IMC.

FFPE sections were placed on Superfrost^®^ Plus slides (Thermo Scientific). Slides were baked at 60 °C for 2 hours in a dry oven and then deparaffinized in fresh xylene before being rehydrated in a graded alcohol series (100%, 96%, 70%; 5 min each). Antigen retrieval was conducted with Tris-EDTA (pH 9) buffer at 95°C in water bath for 30 min. Following cooling, slides were blocked with 3% BSA for 30 min at room temperature. Samples were then stained with the antibody cocktail overnight at 4°C in a hydration chamber. After washing with TBS, slides were incubated with Intercalator-Ir (Standard BioTools) for DNA detection. Finally, slides were washed with TBS and briefly with water, and then air dried. Images were acquired using a Hyperion Imaging System (Standard BioTools) at a laser frequency of 200 Hz. For each sample, regions of interest (ROI) were defined based on the H&E staining for the acquisition of inflammatory areas.

#### Image analysis

##### Tissue and cell segmentation

IMC images were visualized as MCD files using the Visiopharm® software. Each marker was reviewed during an image quality control. Tissue and cell segmentation were then performed using pre-trained deep-learning algorithms based on CD138 and DNA-Iridium channels respectively. All results, multichannel images, and segmentation masks were then exported as standard .tif and .tsv raw data files for subsequent analysis.

##### Phenotyping and cell quantification

Supervised analyses were performed using the OMIC platform (Dotmatics) on biomarkers intensities. Following an initial step of gating on DNA1 and DNA2 to select cells and exclude debris, sequential gating was used to define epithelial cells (CD45-CD138+), CD4+ cells (CD138-CD45+CD3+CD4+), CD8+ T-cells (CD138-CD45+CD3+CD8+), macrophages (CD138-CD45+CD68+), B cells (CD138-CD45+CD20+), naïve B cells (CD138-CD45+CD20+IgD+CD27-), unswitched memory B cells (CD138-CD45+CD20+IgD+CD27+), switched memory B cells (CD138-CD45+CD20+IgD-CD27+), double negative cells (CD138-CD45+CD20+IgD-CD27-), and plasmablasts/plasma cells (CD138+CD45+ CD20+). Percentages and cell densities of immune cells were compared between patient groups by using the Welch’s t test (GraphPad Prism 10 software).

##### Spatial analysis

The spatial analysis pipeline used Phenoplex software (Visiopharm®) for phenotyping. The analysis workflow involved setting thresholds for positivity of each marker through a visual assessment to identify positive and negative cells. Once phenotypes were established for each cell, a hot spot algorithm on the density of the CD20+ cells was designed to visualize B cell distribution. Subsequently, neighborhood matrices were generated using the Python Matplotlib library.

## RESULTS

### Clinical features of lichen planus occurring in patients receiving anti-PD-1 therapy

Between January 2020 and December 2022, six patients who received anti-PD-1 therapy (three with lung carcinoma and three with melanoma) developed lichen planus lesions (Table 2). The median age at the occurrence of the lesions was 71.5 (range 55-77) years with a female prevalence. The time to onset of lichen planus lesions ranged from 4 to 178 (median 46) weeks. The majority of patients developed severe cutaneous eruptions (CTCAE of grade 3 or 4). In parallel, 7 patients with non-drug related LP were included with a median age of 55 (range 39-63) years. All LP patients developed cutaneous eruptions.

**Table 2.** Clinical characteristics of lichen planus patients.

|  |  | ICI-LP (n=6) | LP (n=7) |
| --- | --- | --- | --- |
| **Age, median years (range)** |  | 71.5 (55-77) | 55 (39-63) |
| **Sex, number** |  |  |  |
|  | Female | 4 | 3 |
|  | Male | 2 | 4 |
| **Cancer type, number** |  |  |  |
|  | Lung carcinoma | 3 | NA |
|  | Melanoma | 3 | NA |
| **Stage of cancer, number** |  |  |  |
|  | I-II | 3 | NA |
|  | III-IV | 3 | NA |
| **ICI treatment, number** |  |  |  |
|  | Pembrolizumab | 6 | NA |
| **Time to onset of lesions,**<br>**median weeks (range)** |  |  |  |
|  |  | 45.8 (4-178) | NA |
|  | Unknown | 2 |  |
| **Subtype, number** |  |  |  |
|  | Cutaneous | 5 | 7 |
|  | Oral | 1 |  |
| **CTCAE grade, number** |  |  |  |
|  | 1-2 | 1 | NA |
|  | 3-4 | 4 | NA |
|  | Unknown | 1 | NA |

### Distinct immune infiltration in ICI-induced lichen planus

To profile the immune microenvironment of lichen planus lesions, we used an IMC panel of twelve metal-tagged antibodies targeting CD3, CD4, CD8, CD20, CD27, CD38, CD45, CD56, CD68, CD138, IgD, and pan-cytokeratin. Serial sections stained with H&E confirmed features of lichenoid interface dermatitis in both group of patients (Figure 1A) and were used to select ROI for IMC acquisition. In total thirty-four ROI were analyzed, fourteen from LP patients and 20 from ICI-LP patients. As shown in Figure 1B, the panel allows the characterization of epithelial cells (CD45-CD138+), CD4+ T-cells (CD45+CD4+), CD8+ T-cells (CD45+CD8+), B cells (CD45+CD20+), and macrophages (CD45+CD68+).

**Figure 1.**
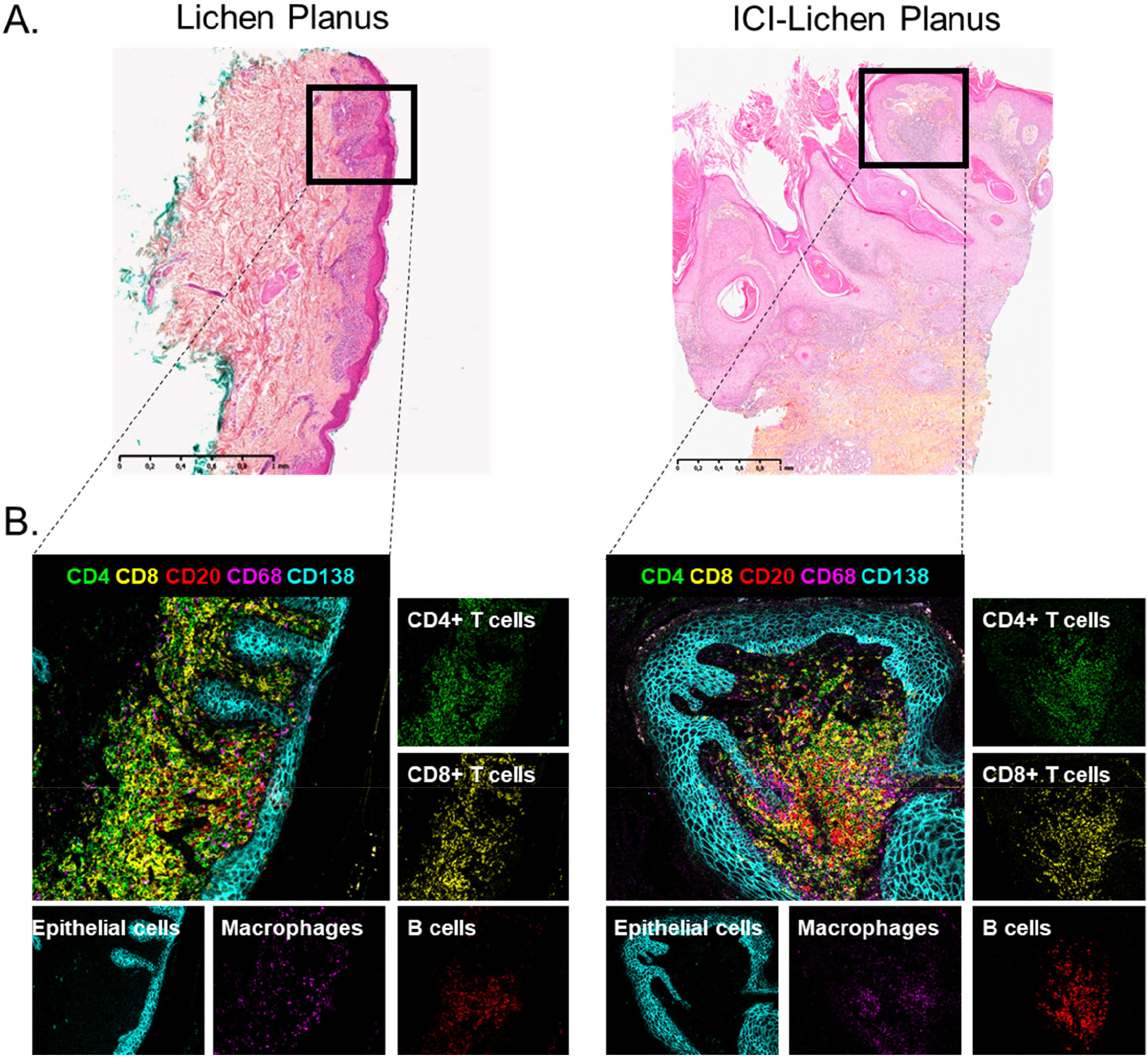
Visualization of the tissue structure microenvironment in single regions of interest in lichen planus by IMC. A. Representative H&E staining of lichen planus and ICI-lichen planus with selected ROI for IMC acquisitions. B. Representative IMC image of lichen planus and ICI-lichen planus showing the overlay and the individual stains of CD4 (green), CD8 (yellow), CD20 (red), CD68 (magenta), and CD138 (cyan).

All images were segmented using a deep-learning algorithm, which initially identified the tissue, and subsequently separated it into dermis and epidermis (Figure 2A). The cells within the tissues were then segmented based on DNA-Iridium channels. Visualization through T-SNE plots revealed that the majority of immune cells were located in the dermis of LP, while epithelial cells were confined to the epidermis (Figure 2B). Using a supervised approach, we found a trend towards a decrease in the density of CD45+ immune cells per mm2 in ICI-LP compared to LP (Figure 2C). Detailed quantification of cell densities revealed that the immune infiltrates were predominantly comprised of T cells (LP 76%, ICI-LP 84%). However, a significant decrease of CD4+ T-cell density was observed in ICI-LP compare to LP. Additionally, cell densities showed also a trend toward decreased CD8+ T-cell density associated with an increase of B cell density. Interestingly, the frequency of CD20+ B cells within CD45+ immune cells was significantly increased in ICI-LP compared to LP (Figure 2D). Altogether, using a high-dimensional spatial characterization by IMC, we highlighted the reduction of T-cell infiltration in ICI-LP compared to LP, concomitantly to the increase of B-cell frequency.

**Figure 2.**
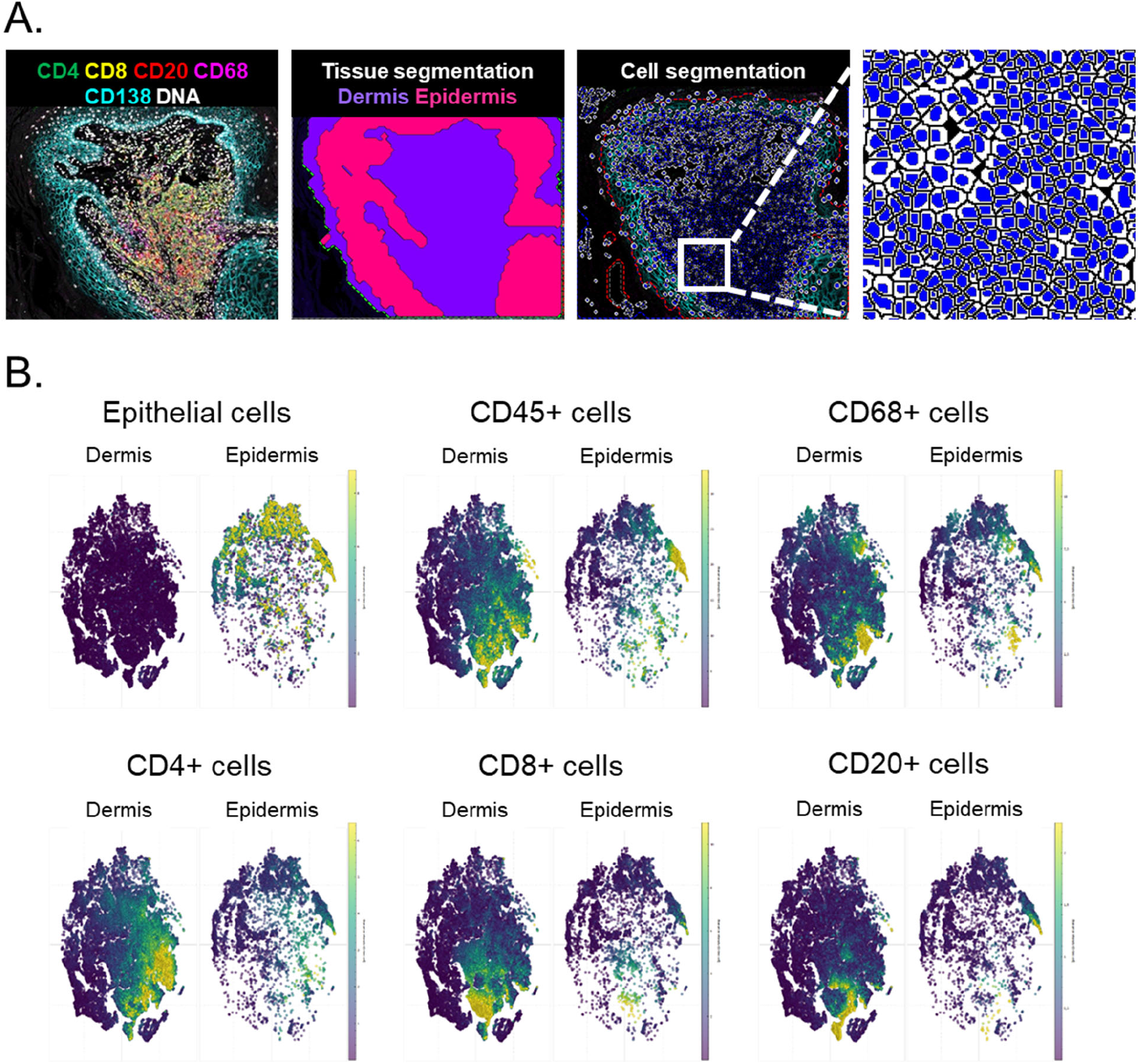
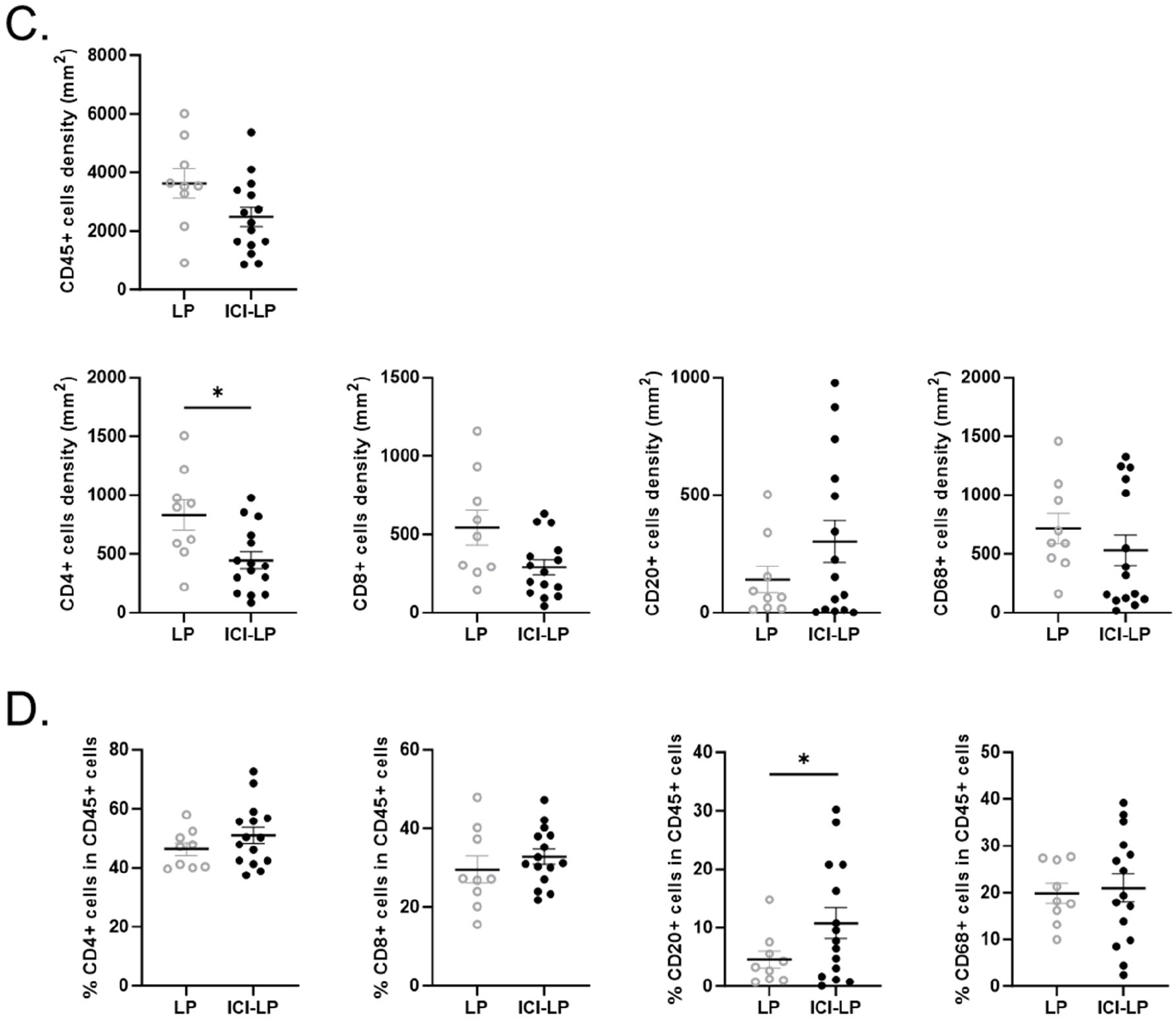
Immune composition of lichen planus. A. Tissue and cell segmentation using Visiopharm® software. B. T-SNE plots displaying the expression of CD138, CD45, CD68, CD4, CD8, and CD20 in both the dermis and epidermis. C. Cell density of immune cells in LP and ICI-LP. D Immune cell subset frequencies in both groups of patients. Welch’s t tests were performed: *P < 0.05. Abbreviations: ICI, immune checkpoint inhibitor; LP, lichen planus.

### B-cell changes in ICI-induced lichen planus

We then analyzed the distribution of B-cell subpopulations based on the differential expression of several differentiation markers.^11^ Thus, five B-cell subsets were defined based on the expression of IgD, CD27, CD138 on CD20+ B cells; naïve B cells (IgD+CD27-), unswitched memory B cells (IgD+CD27+), switched memory B cells (IgD-CD27+), double negative cells (IgD-CD27-), and plasmablasts/plasma cells (CD138+CD45+) (Figure 3A). The majority of B cells were double negative B cells in both groups of patients (Figure 3B). However, a significant decrease in double negative B cells was observed in ICI-LP compared to LP, concomitantly to an increase in naive B cells. For plasmablasts/plasma cells, no significant difference was found. Taken together, these data show alteration in B cell subset distribution in ICI-LP compared to LP.

**Figure 3.**
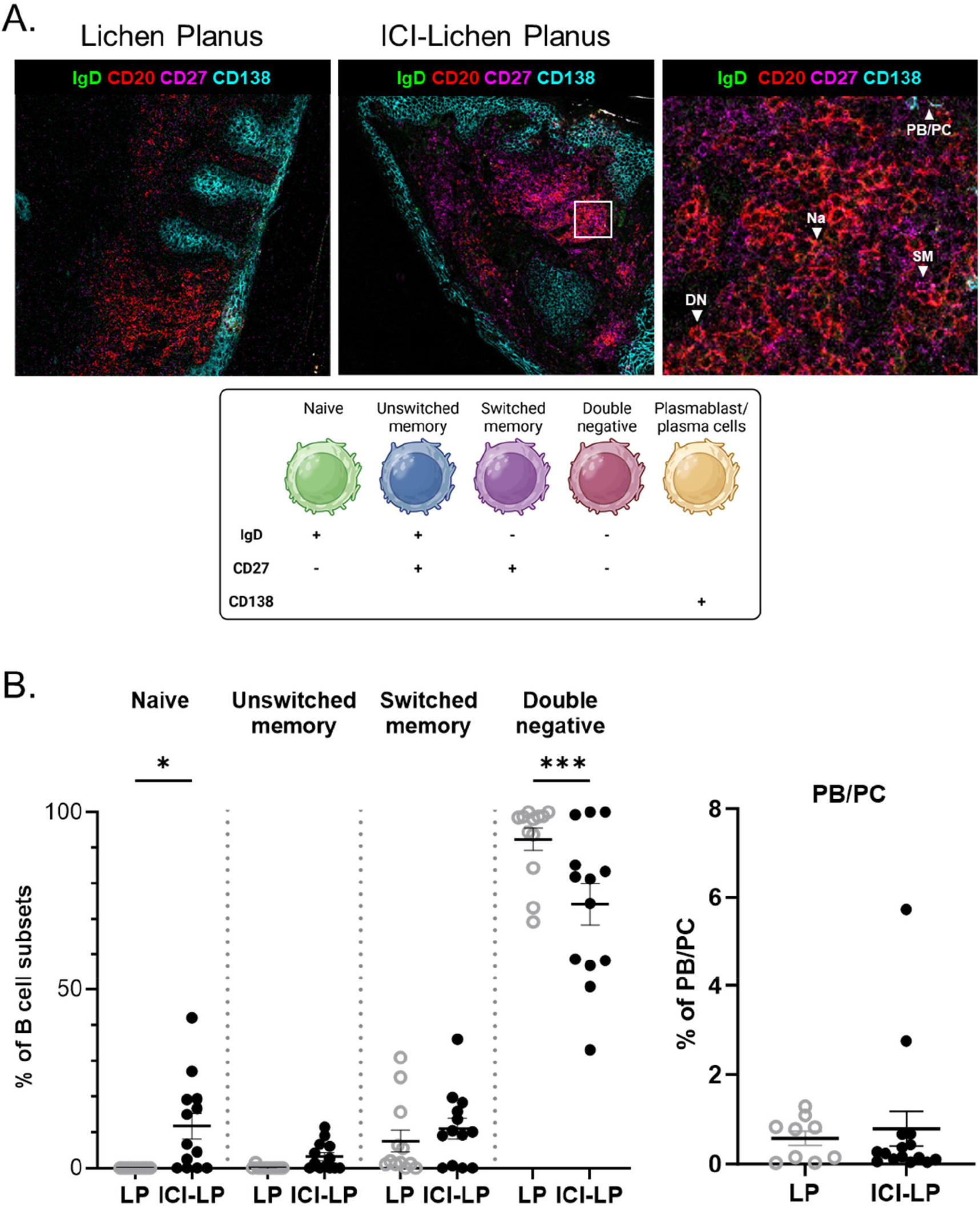
Phenotypic characterization of infiltrating B cells in lichen planus. A. Representative IMC images of lichen planus and ICI-lichen planus showing IgD (green), CD38 (yellow), CD20 (red), CD27 (magenta), and CD138 (cyan). B. Frequency of B-cell subsets in LP and ICI-LP. ANOVA (One Way) Sidak test and welch’s t tests were performed: *P < 0.05 and ***P < 0.001. Abbreviations: ICI, immune checkpoint inhibitor; LP, lichen planus.

### B cell are organized in aggregates close to T cells

To analyze the spatial distribution of B cells, we first visualized cell phenotyping and B cell hot spots using a heatmap based on the density of the CD20+ cells (Figure 4A). In ICI-LP, we observed B-cell aggregates surrounded by other immune cells, whereas B cells were more sparsely distributed in the immune infiltration of LP. Neighborhood matrices revealed that B cells resided next to CD8+ T cells (LP 37%, ICI-LP 38%) and were in avoidance with macrophages and epithelial cells in both group of patients (Figure 4B). Interestingly, ICI-LP showed a higher frequency of CD4+ T cells next to B cells compared to LP (LP 10%, ICI-LP 17%). We also observed a decrease of CD8+ T cells frequency in proximity of epithelial cells in ICI-LP (LP 33%, ICI-LP 26%). These findings revealed that infiltrating B cells are organized in aggregates closed to T cells in ICI-LP, suggesting a potential role in the local adaptive immune response.

**Figure 4:**
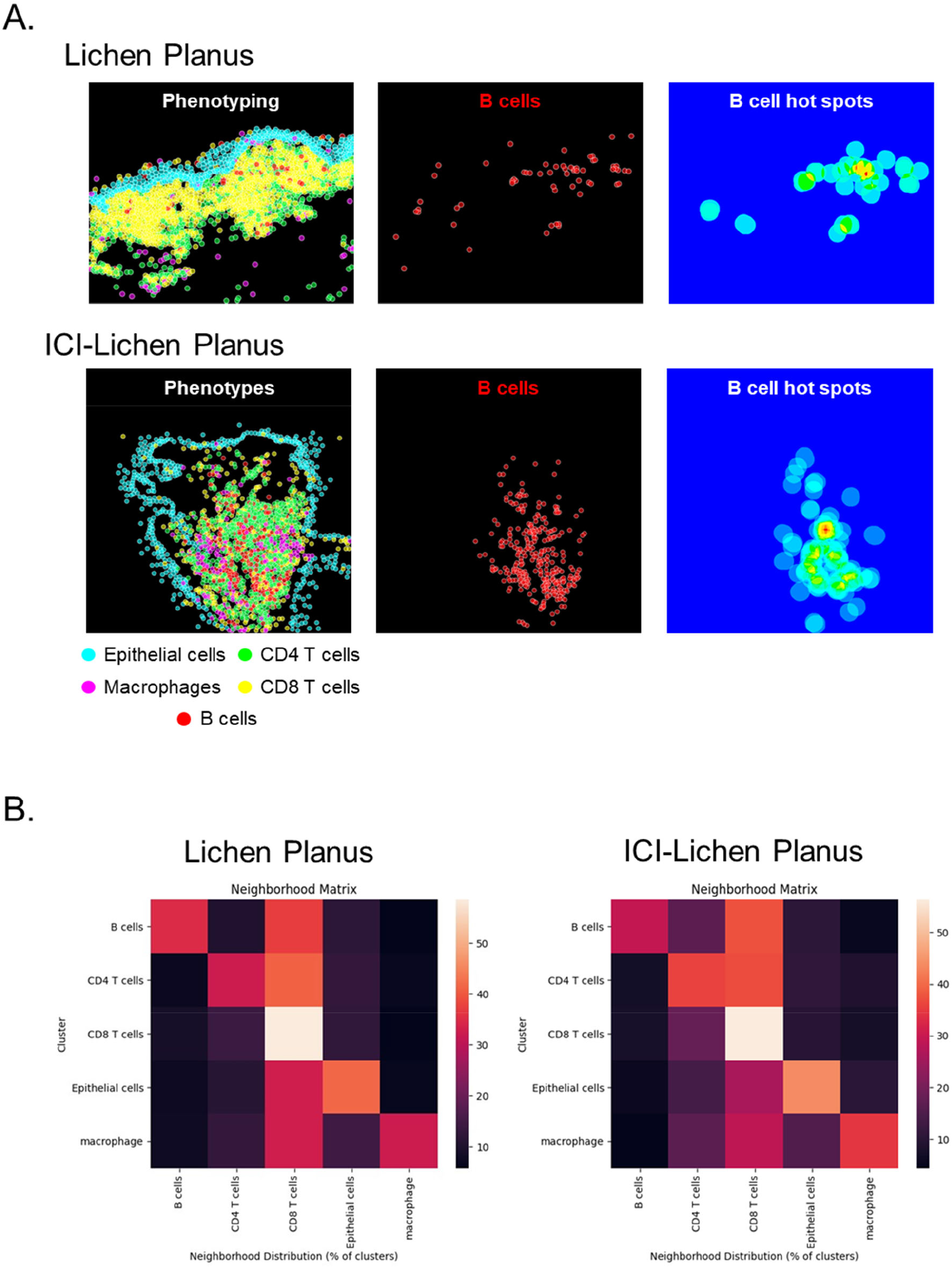
Spatial analysis reveals B-cell aggregates in closed proximity with T cells. A. Representative images of phenotype classification and heatmaps based on the density of the CD20+ cells in LP and ICI-LP. B. Heatmaps showing the distribution of neighborhood clusters.

## DISCUSSION

Here, we provided an in depth analysis of immune infiltration in lichenoid reactions occurring in patients undergoing anti-PD-1 therapy using IMC. This approach allowed to identify the spatial organization of immune cells within the skin. Our data revealed for the first time distinct pattern of immune infiltrates in ICI-LP compared to LP. We observed a simultaneous decrease in T cells and an increase in B cells in ICI-LP. B-cell infiltrates in ICI-LP exhibited a decrease in double negative B cells in association with an increase in naïve B cells. These B cells were organized in aggregates close to CD4+ and CD8+ T cells. Overall, our data support the concept that B cells play a role in the physiopathology of c-irAE.

Cutaneous complications are among the most prevalent irAE and occur in approximately 30% to 60% of patients receiving anti-PD-1 antibodies.^12^ Although c-irAE are among the earliest complications to develop within the first weeks to months after treatment initiation, the onset of these complications differs depending on the type of skin rash. The onset of lichenoid reactions occurs later than psoriasiform rash and maculopapular rash, ranging from 6 to 12 weeks.^13^ Lichenoid eruptions are more frequent in patients receiving anti-PD-1/PD-L1 antibodies than in those receiving anti-CTLA-4 antibodies.^14^ In our study, all patients received anti-PD-1 therapy (pembrolizumab), and the majority experienced grade 3 toxicities. The high prevalence of severe grades of ICI-LP in our cohort may be explained by the fact that a skin biopsy was not performed in patients with low-grade cutaneous irAE. Moreover, the time to onset ranged from 4 to 178 weeks, with a median at 45 weeks. A recent study found that in a real-world clinical setting, cutaneous diagnoses are made later than reported in clinical trials.^15^ The median time to the first cutaneous irAE in this cohort of 8637 patients receiving ICI therapy was 16 weeks for any type of cutaneous manifestations. In addition, they demonstrated that lichen planus diagnoses are made later with a median time of onset of 30 weeks (17-50 weeks). These findings in a real-world setting suggest that patients are referred later to dermatologists due to a lack of urgency or access outside clinical trials, which may explain their progression to a higher grade of toxicity.

Another notable finding is that despite the decrease in T cells in ICI-LP compared to LP, lichenoid eruptions remained dominated by T-cell infiltrates. While the pathophysiology of c-irAE is mainly unknown, several studies reported T-cell mediated mechanism.^7-9^ A mice model, which enables *de novo* induction of skin-specific expression of T cells antigens in the epidermis, demonstrated that ICI treatment led to localized cutaneous disease.^16^ Skin-infiltrating antigen-specific CD8+ T cells were characterized by the expression of genes associated with T-cell activation, effector function and immune checkpoints. Moreover, they confirmed the infiltration of CD8+ T cells with effector capacity in punch biopsies from two patients with ICI-LP. These findings support that PD-1 maintains local CD8+ T cells tolerance and prevents skin pathology.

Additionally, to these previous studies, we described an increased frequency of infiltrating B cells in ICI-LP compared to LP. In line with these results, Tsukamoto *et al*. demonstrated an increase in B cells organized in dense aggregates, resembling tertiary lymphoid structures (TLS), in the lungs and liver of tumor-bearing aged mice treated with ICI, which were characterized by multiorgan toxicities.^17^ They demonstrated that ICI treatment upregulated CXCL13 expression, which is involved in the accumulation of B cells in organs affected by irAE and in the formation of TLS. In addition, the study reported that CXCR5, the receptor of CXCL13, was expressed only on B cells, not on T cells, in the organs of in ICI-treated aged mice, suggesting the involvement of CXCL13 in B cell migration within TLS. Our data show, for the first time to our knowledge, the organization of B cells in aggregates in human organs affected by irAE. Although the presence of TLS-related B cells has been linked to improved survival and sensitivity to ICI therapy in several cancers,^18,19^**^-^**^21^ a detailed analysis of B cells in irAE will provide a better understanding to decouple antitumor efficacy from irAE.

## CONCLUSION

Our study reveals significant changes in the immune infiltrate of ICI-LP compared to LP, with a concurrent decrease of T-cell infiltration and an increase in B cells organized in aggregates. These findings suggest a distinct physiopathology between these two cutaneous manifestations.

## ACKNOWLEDGEMENTS

The authors acknowledge the flow cytometry core facility Hyperion (Brest, France) for its technical assistance and the European subsidy programme FEDER Progos RU 000950.

